# A triple fluorescent marker for live imaging of plant cell morphogenesis

**DOI:** 10.64898/2026.08.20.745988

**Authors:** Zoé Bomsel, Coralie Goncalves, Aloïse Ducamp, Loris Caillat-Miousse, Bérengère Dalmais, Katia Belcram, Chie Kodera, Claire Lionnet, Camila Goldy, Samantha Moulin, Marie-Cécile Caillaud, David Bouchez, Martine Pastuglia, Magalie Uyttewaal

## Abstract

Live imaging of plant subcellular structures is key to deciphering the spatiotemporal bases of cellular processes, and their functional impact on growth and morphogenesis at various biological scales. Live imaging of plant cells essentially relies on expression of fluorescent markers labeling cells or subcellular structures of interest. Simultaneous multi-channel imaging of several markers is still not routine practice in plant cell biology, owing to issues linked to genetic or spectral compatibility of markers, differences in expression levels, silencing, toxicity, etc. Here we designed a three-color marker in *Arabidopsis thaliana* and *Capsella rubella*, enabling high-resolution live imaging of plant morphogenesis, including labeling of the cell membrane, the nucleus and the microtubule cytoskeleton. Detection of MT arrays involved the development of a MAP4-MBD-based microtubule marker optimized for plant cells. The three-color marker allows visualization of the three-dimensional organization and dynamics of plant microtubules within the intracellular space with unprecedented precision, in various organs including the root and shoot meristems, the leaf, anther, and gynoecium. Our results demonstrate the potential of such single-construct strategy for cell biology studies in plants.

## Introduction

Live imaging of plant subcellular structures such as the cell wall, the nucleus, the membranes, vesicles, or the cytoskeleton is instrumental to the study of spatiotemporal bases of cellular processes, and of their functional impact on growth and morphogenesis at various biological scales. Advances in plant cell imaging strongly depend upon the continuous development of a variety of fluorescent probes and fluorescent proteins (FPs) (e.g. [1]), on rapid progresses in confocal and super-resolution microscopy, as well as recent ones in expansion microscopy [2–4]. A comprehensive toolbox for microscopic analysis of plant cells is readily available to plant biologists, and continues to expand, allowing visualization of specific tissues (e.g. [5]), cell cycle stages [6], and a variety of subcellular structures including the cytoskeleton [1,7], the nucleus [8], or membrane domains [9], to name a few.

However, in practice, FP markers suffer from a number of drawbacks that limit their potential applications [10]. First, marker expression is highly variable among transgenic lines, possibly leading to artifactual localization or phenotypic alterations when marker expression is under or above a certain threshold. Although this strategy is not recommended under current standards, many first-generation plant cell markers were produced under the control of strong constitutive promoters such as the 35S promoter from the Cauliflower Mosaic Virus, leading to overexpression well above physiological levels. Overexpression of FP fusions, in turn, often leads to cytotoxicity and impairment of growth and development, from the cellular to the whole plant level [11]. In the same lines, mis- and over-expression frequently induce silencing in plant cells, producing highly variable and patchy patterns of marker expression. Some constructs may even inherently induce silencing, sometimes precluding selection of a single marker line devoid of silencing defects. Moreover, above a given level of expression, most (sub-)cellular markers display at least some level of toxicity, which can hamper their use in practice.

An archetypal example in the field of plant cell biology is fluorescent fusions with the microtubule-binding domain (MBD) of mammalian MAP4 [12]. Since the original publication of the first p35S-GFP-MAP4 MBD marker in 1998 [13], FP-MBD fusions have been widely used as live microtubule (MT) markers in plant cells, owing to their excellent signal-to-noise ratio both in interphase and mitosis, low cytoplasmic background, and absence of any homologue in the plant kingdom with which they could interfere or compete. However, strong expression levels of MAP4-MBD constructs in plant cells can lead to mild to severe growth defects [14], slower proliferation of cell suspensions [15] or mild organ twisting [16], due to the intrinsic MT bundling activity of the MAP4-MBD [17]. MT bundling refers to individual microtubules being laterally cross-linked, to form closely aligned arrays of (anti-)parallel or mixed polarity. *In vivo*, bundling is mediated by specific MAPs that connect adjacent microtubules and play essential functions in eukaryotic cells including in plants (e.g. [18]). The use of lines heterozygote at the MT marker locus, together with less potent promoters [19] and stringent selection of transgenic lines for the absence of phenotype can often alleviate these caveats, but, in general, to the cost of signal brightness. Alternative MT markers such as FP fusions with α- and β- tubulins are also widely used in plant cell research, but often display higher cytoplasmic backgrounds than MAP4-based markers, and are not devoid either of silencing or toxicity issues, especially when expressed at high levels [14,20].

As a consequence of such drawbacks associated with expression of FP fusion markers in plant cells, the obtention of stable marker lines generally necessitates heavy selection procedures and careful examination of potential phenotypes, in order to fine-tune the expression level of the marker(s) without compromising either the quality of the signal or the dynamics of the structure under study.

The task becomes even more challenging when it comes to multi-channel live imaging, where several markers are to be recorded simultaneously. In this case, optimizing the expression levels and harmlessness of two, three or more markers simultaneously, while keeping them compatible in terms of stability, brightness, absence of spectral overlap, or genetic position can become a daunting challenge. Optimizing each marker line individually in a wild-type background, then combining the markers by crossing to finally produce a homozygous wild-type reference line remains a preferred option in most cases. This reference line, in turn, can be crossed with other lines to transfer the markers into any genetic background. However, genetically unlinked markers are difficult to work with and, depending on the number of markers and mutations in segregation, obtention of a fully homozygous line can be time consuming and may necessitate several generations, even provided that all markers and mutations segregate independently. Moreover, the accumulation of transgenes and selectable markers in multiple mutant lines may strongly increase the propensity for silencing of transgenic loci.

In order to –at least partially– alleviate these limitations, we set out to optimize a single-locus, triple marker construct designed to simultaneously label the cell membrane, the nucleus and the microtubule cytoskeleton by three-color live imaging. We first designed a new microtubule marker based on a truncated MAP4-MBD isoform previously shown to strongly reduce the MT bundling activity of the MAP4-MBD [21] *in vitro*. We developed a MT bundling analysis pipeline from confocal images and showed that the new 3R-MAP4-MBD marker significantly reduced the bundling propensity of the original 5R version in stably transformed *Arabidopsis* cotyledon cells. Using the GoldenBraid cloning strategy [22], we produced a series of individual markers that we grouped into a transformation-ready triple marker vector. Lines expressing this triple marker were selected in both *Arabidopsis thaliana* (wild-type and mutants) and *Capsella rubella.* This triple marker notably allows exploration of the dynamic organization of the microtubule cytoskeleton within the 3D space of the plant cell, providing crisp views of both cortical and intra-cytoplasmic MTs and their connections with the nuclear envelope. High resolution 3-channel imaging of several organs including the root and shoot meristems, young leaves, the hypocotyl or the gynoecium demonstrated the broad application potential of this marker for developmental studies in plants.

## Materials and methods

### Molecular cloning, bacterial and plant transformation

The GoldenBraid cloning system [23] was used for all constructs. GoldenBraid parts were either available from Addgene (https://www.addgene.org/) or domesticated in pUPD2. Vectors were assembled using *Bsa* I or *Bsm* BI type II restriction enzymes, according to the GoldenBraid 2.0 protocol [24] (https://goldenbraidpro.com/). Plasmids used to create transcription units (TU) or multigene assemblies were pDGB3 series. Binary vectors were transformed into electrocompetent *Agrobacterium tumefaciens* C58C1-pMP90 [25] by electroporation. Transformed *A. tumefaciens* were re-suspended in a 2 mM MgCl_2_, 5% (w/v) sucrose, 0.02% Silwet solution and infiltrated into *A. thaliana* by floral dip [26].

DNA from the MNM line was prepared for whole-genome sequencing using the NucleoSpin Plant II kit (Macherey Nagel). DNA extraction was performed on 15-day-old plants grown *in vitro*. Library preparations and Hi-Seq Illumina sequencing (2 × 150 bp paired ends) were performed by Eurofins Genomics. The script used for aligning reads against the reference genome is described in [27].

Oligonucleotides, transcription units, and DNA sequence of the MNM marker transformation vector are available as supplementary materials (**Supplemental Files 1 to 3**).

### Plant materials and cultivation

All *Arabidopsis thaliana* plants used in this study were in the Columbia-0 (Col0) background. The *katanin* P60 mutant allele is *bot1-6* [28]. The *Capsella rubella* ecotype was Cr22.5 [29].

Greenhouse phenotyping was performed under long-day conditions (16 h light at 23°C and 8 h dark at 17°C). Seeds were stratified for 2 days at 4°C in 0.1% (w/v) agar before being sown directly onto soil in 7 × 7 × 6.5 cm square plastic pots arranged in rigid trays. Genotypes were randomly assigned to positions within the trays to minimize positional effects. Plants were imaged using a Canon EOS 500D camera mounted on a fixed imaging platform. Rosettes from greenhouse plants were segmented by thresholding and their area was quantified using the MorpholibJ plugin in ImageJ/Fiji [30].

For *in vitro* root phenotyping, seeds were sown on square Petri dishes containing half-strength Murashige and Skoog (½ MS) medium supplemented with 1% (w/v) sucrose and 0.8% (w/v) Phytoblend. Seeds were placed along the upper third of the agar surface to allow unrestricted root growth during vertical cultivation. Following a 2-day stratification period at 4°C, plates were transferred to a growth chamber under long-day conditions (16 h light at 21°C and 8 h dark at 18°C). Root systems were scanned at 2 and 7 days after transfer using an Epson Perfection V850 Pro flatbed scanner. Primary root length was measured at 2 and 7 days after germination using the NeuronJ plugin for ImageJ [31]. Root growth was then calculated as the increase in root length between D2 and D7, thereby correcting initial root length variability resulting from asynchronous germination.

### Confocal imaging and image processing

Confocal images of root tips were acquired with the following set-ups: 1) SP8 confocal microscope (Leica) equipped with a 40x PlanApochromat objective (NA 1.30, oil immersion) or 2) LSM980 Airyscan 2 confocal microscope (Zeiss) equipped with a 40x C-Apochromat objective (NA 1.2 water immersion) or 3) inverted Zeiss microscope (AxioObserver Z1) equipped with a spinning-disk module (CSU-W1-T3, Yokogawa) and a CameraPrime 95B (Photometrics) using a 63x PlanApochromat objective (NA 1.4, oil immersion). The images of shoot apical meristems, anthers and cotyledon pavement cells were acquired using a Leica upright SP8 confocal microscope equipped with a 25x HC Fluotar (NA 0.95 W, water immersion) lens. The images of hypocotyls and petals were acquired using an SP8 confocal microscope (Leica) equipped with a 40x PlanApochromat objective (NA 1.30, oil immersion). Images of interphasic cortical microtubules for bundling analysis were acquired using a Leica Stellaris 8 confocal microscopy equipped with a 40x PlanApochromat objective (NA 1.3)).

Calcofluor staining and imaging of *Arabidopsis* roots were performed as described previously [32].

All image processing was done in the Fiji /ImageJ software [33]. Where applicable, the SurfCut macro [34] was used to project the signal at chosen distances relative to the surface of the cell (e.g. between 0 and 2 μm for a cortical signal, between 2 and 4 μm for a median section).

An original MT bundling analysis pipeline was designed to compare the MT bundling activity of the new 3R- marker in comparison to the original 5R-MBD version. The conceptual basis of the pipeline stems from the observation that the maximum gray value of MT bundles is proportional to the number of MTs within the bundle, provided the MT signal is not saturated (**Fig. 3 A-D**). For details on the pipeline and its use, see **Supplemental Files 4 and 5**. Briefly, a stack of interphasic cortical microtubules was first denoised using a 3D median filter, and a maximum projection was generated. The microtubule signal was then segmented by binarization, followed by an erosion/dilation cycle to eliminate small background particles. This binary mask was then applied to the original maximum projection, resulting in an image in which most non-MT cytoplasmic background was suppressed. Next, the image was cropped in order to keep the central part of the projection, encompassing the flattest part of the sample. Visual inspection of the cropped image and of its histogram then allowed to estimate the average maximum gray value of single microtubules. This value was then used to divide the image, resulting in an image in which pixels of value 1 represent single microtubules (MT1), pixels of value 2 represent bundles of two microtubules (MT2), and so on. Simple pixel counting then allowed approximation of MT bundling rate (BR) by the formula:

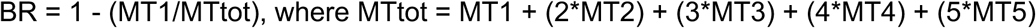

The whole procedure was semi-automatized and coded as a Fiji/ImageJ macro (**Supplemental File 4**) that stops for user input and validation when necessary. A detailed user manual is provided (**Supplemental File 5**). To avoid potential biases, images of cells expressing the 3R- and 5R-MBD markers were anonymized and randomized prior to analysis.

## Results and discussion

### A new MAP4-MBD based microtubule marker

Mammalian MAP4 belongs to the Tau family of microtubule-associated proteins (MAPs). This family of MAPs plays on microtubule stability and regulates microtubule-based motility in vertebrates. They include a conserved C-terminal microtubule-binding domain composed of three (3R) to five (5R) tubulin-binding repeats and an N-terminal projection domain that is specific to each member of the family [35]. For Tau, MAP2 and MAP4, splice variants harboring different numbers of repeats are produced in a tissue-specific manner, the number of tubulin-binding repeats having a significant effect on the activities of these MAPs [35]. MAP4 is abundant in non-neuronal cells, and contrary to Tau and MAP2, does not induce strong bundling of MTs in its full-length form [17]. It has been shown that this difference stems from the N-terminal projection domain of MAP4, which inhibits the intrinsic bundling activity of the C-terminal MT-binding domain [17]. Interestingly, the bundling activity of the MAP4-MBD strongly depends on the number of tubulin-binding repeats [21]. In an *in vitro* study, 3R, 4R and 5R versions of the bovine MAP4-MBD were tested for their ability to induce lateral association of MTs. The 5R version induced strong bundling, with less than 10% of single MTs, and 90% involved in bundles of two, three or four MTs. On the contrary, with the 3R version, 80% of the microtubules were single, and only 20% were bound to another microtubule, while less than 1% were engaged in larger bundles of three to four microtubules. The 4R version displayed intermediate bundling rates [21].

As the original MAP4-MBD construct used as a MT marker in plants [13] as well as all its derivatives to date (eg [19]) are 5R versions (**Fig. 1A**), we designed a new, 3R version of this marker and tested its ability to label plant MTs while reducing the MT-bundling propensity classically affecting the 5R versions [14–16]. To this end, we deleted a 69-amino-acid region encompassing repeats R2 and R3 in the original mouse MAP4-MBD version in order to produce a strict equivalent of a mammalian 3R-MBD isoform, retaining only the R1, R4 and R5 repeats (**Fig. 1A**).

**Figure 1.**
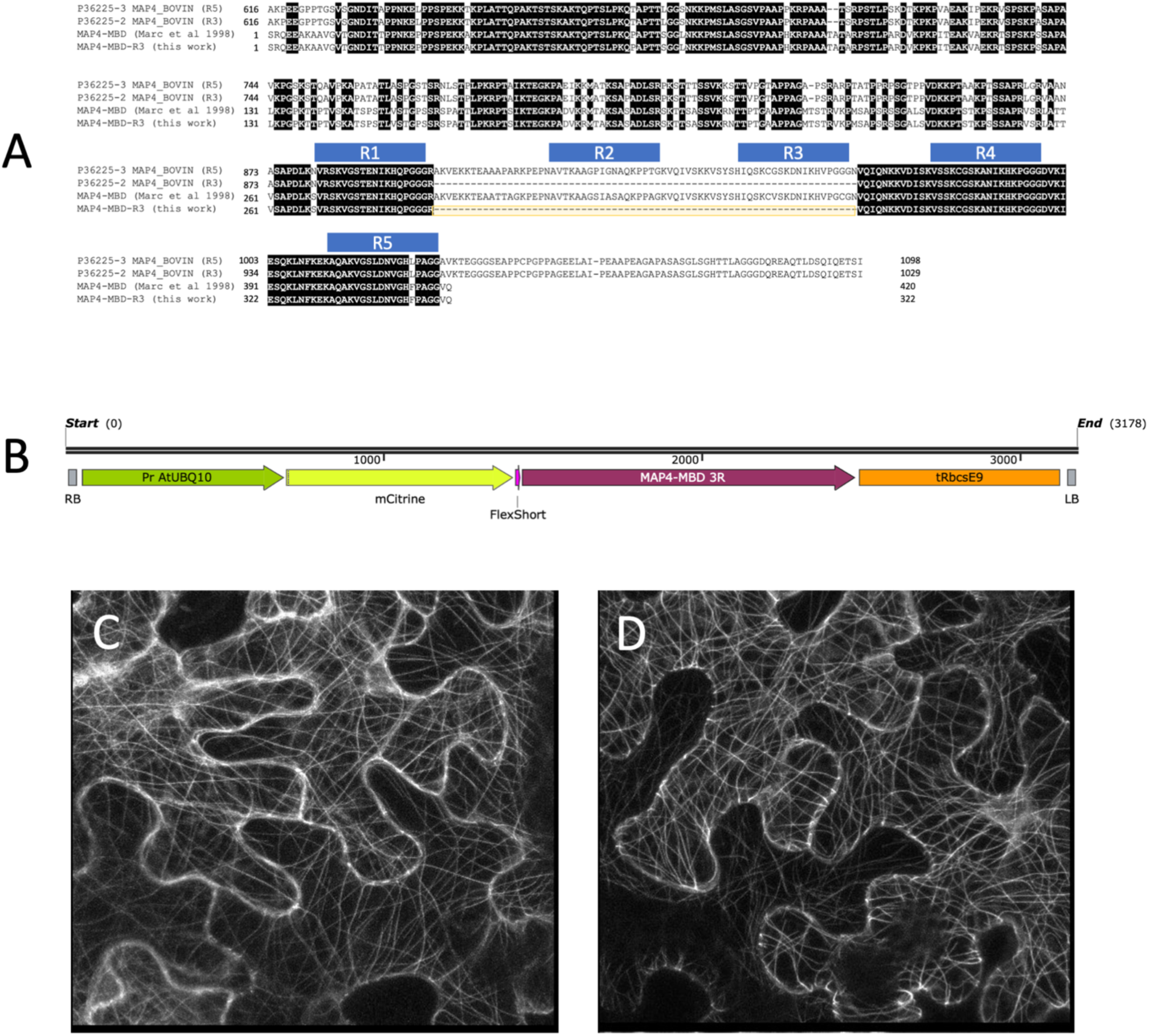
Design of a new 3R-MBD microtubule marker for plant cells. **(A)** Partial protein alignment of the 5R and 3R versions of bovine MAP4 (Uniprot P36225), together with the original MAP4-MBD construct [13] and its 3R derivative (this work). The tubulin-binding repeats are highlighted in blue. The R2-R3 region deleted in 3R isoforms and our 3R construct (69 residues) is highlighted in yellow. **(B)** The T-DNA of the pDGB3_Alpha2R::P_UB10_::mCitrin::FS::3R-MBD::T_RbcsE9_ vector consists of a promoter fragment from the *Arabidopsis* At4g05320/UB10 gene encoding polyubiquitin 10, which comprises 636 bp upstream of the ATG. The UB10 drives expression of an mCitrin-FS-3R-MBD translational fusion, in which FS stands for flexible short linker (GlyGlyGlyGlySer). A 631 bp terminator fragment from the RbcsE9 gene from pea is placed downstream of the fusion. Total size of the construct is ∼3 kb. Comparison of 5R-MBD **(C)** and 3R-MBD **(D)** mCitrine fusions upon transient expression in *Nicotiana benthamiana* leaf cells. Constructs are P_UBI_::mCitrine::5R- MBD::T_NOS_ in **(B)** and P_UBI_::mCitrine::3R-MBD::T_NOS_ in **(C)**, differing only by the number of tubulin-binding repeats.

In order to assess the functionality of the 3R-MBD version in plant cells, a P_UB10_::mCitrine::3R- MBD::T_NOS_ fusion construct was assembled in the GoldenBraid destination vector pDGB3-α2R. This vector was introduced into *Agrobacterium*, and first used in transient expression assays by infiltration of *Nicotiana benthamiana* leaves, in comparison to the original 5R version in the very same configuration. In these non-cycling leaf epidermal cells, the whole cortical microtubule array was clearly labeled, and the signal-to-noise ratio appeared comparable to the original 5R construct, showcasing the functionality of the marker (**Fig. 1C, 1D**) and confirming that the 3R and 5R versions of the MAP4-MBD are quite similar in terms of MT affinity, as shown previously [21].

Comparing N-terminal and C-terminal mCitrine fusions with the 3R-MBD upon transient expression in *N. benthamiana* cells and in *Arabidopsis* stably transformed lines did not reveal any significant difference in terms of MT labeling (not shown). Likewise, linkers of variable size and flexibility –two rigid and two flexible– were placed between the mCitrine fluorescent marker and the 3R-MBD. Their effect on marker brightness and stability was assessed by transient transformation of *N. benthamiana* leaf pavement cells and in *Arabidopsis* stably transformed lines, and did not reveal any observable difference (not shown). We thus chose to pursue with a construct harboring the mCitrine at the N-terminus and the microtubule marker in C-terminal position, as in the original GFP-5R-MBD construct [13], a short flexible linker (FS) connecting the two parts (**Fig. 1B**).

Next, we compared the 3R-MBD and 5R-MBD fusions in stable transgenic lines of *Arabidopsis thaliana*. To get a more quantitative view, we compared the 5R and 3R versions driven either by the P_35S_ or the P_UB10_ promoter (**Table 1**). A total of 14 to 22 independent transgenic lines were analyzed per construct. For each line, signal brightness was assessed in 3-4 roots from 3-day-old plantlets grown *in vitro*. Signal intensity was approximated as inversely proportional to the photodetector gain required to obtain a clear, standard MT image, all other acquisition parameters being kept constant.

**Table 1.** Comparison of signal intensity in various MT markers. A comparison was made between P_35S_ *vs*. P_UB10_ -driven constructs, and between 3R and 5R versions of the MAP4-MBD. The number of transgenic lines analyzed for each construct is shown. Root images (>3 per line) were acquired in the same exact conditions on a Leica SP5 confocal microscope (514 nm ray at 20%, hybrid detector). Only gain was corrected to visually adjust all signals to the same brightness. Signal strength is approximated here as inversely proportional to the level of gain necessary to adjust an image.

| Construct | #<br>*lines* | Detector Gain |  |
| --- | --- | --- | --- |
|  |  | 10-100 | 100-300 |
| P<sub>35S</sub>::mCit-FS-5R-MBD | 18 | 94% | 6% |
| P<sub>35S</sub>::mCit-FS-3R-MBD | 14 | 72% | 28% |
| P<sub>UB10</sub>::mCit-FS-5R-MBD | 20 | 25% | 75% |
| P<sub>UB10</sub>::mCit-FS-3R-MBD | 22 | 18% | 82% |

The proportion of lines in low or high gain classes (**Table 1**) confirmed that P_35S_-driven constructs were expressed at very high levels, mostly necessitating gain values of less than 100, while P_UB10_-driven constructs yielded lower signals, with a majority of lines requiring gains in the range of 100 to 300 at 30% laser power. This confirmed that the P_UB10_ promoter is of moderate activity [36] compared to the P_35S_ one, and clearly a better choice to avoid toxicity or silencing issues induced by overexpression of MBD fusions. This analysis also corroborates that the 5R- and 3R- versions of the MBD yield comparable signals, although the 5R, on average, consistently produces slightly brighter signal than its 3R counterpart.

From this, we concluded that the 3R-MBD, although somewhat less bright, yields a clear and homogeneous MT signal and represents a good alternative to the original 5R-MBD construct.

Next, we set out to evaluate the microtubule bundling activity of the 3R- marker as compared to the original 5R-MBD. Despite their biological relevance, MT bundles are not possible to resolve from single MTs in diffraction-limited microscopy systems [37], and the resolution of individual MTs within bundles is not accessible even with super-resolution imaging [38,39]. In addition, to the best of our knowledge, no standard approach exists for the estimation of MT bundling from fluorescence microscopy images. Efforts to develop such methodology are surprisingly scarce in the literature and mostly aimed at *in vitro* experiments [40,41].

In high- or super-resolution low noise images, it is demonstrated that MT signal intensity is proportional to the number of MT(s) composing the fiber [37,38], allowing determination and counting of filaments corresponding to one, two, or more individual MTs based on cross-profile analysis and area under the curve calculations [37]. Here we show that the peak value of the signal in regular, live confocal images is also proportional to the number of MT within the bundle, and that this property can be used to estimate the MT bundling rate in a region of interest (**Fig. 2A-D**). To that end, we developed an original pipeline for assessing MT bundling in regular scanning confocal microscope images of cells expressing an MT fluorescent reporter. The method presented here has a few identified biases due for example to the bell shape of the MT signal intensity (**Fig. 2C**) that tends to lead to underestimate bundles, since border pixels are classified as lower-order bundles. On the contrary, MT crossings are counted as higher-order bundles, thereby overestimating bundle counts (**Fig. 2D**). To improve the precision of the method, more advanced approaches could be deployed like masking the image with a skeletonized mask, or using an area under the curve integration method as in [37]. Nevertheless, while not intended to produce a precise and absolute bundling rate, the method allows to compare the intensity of bundling between conditions or markers acquired in homogeneous imaging conditions.

**Figure 2.**
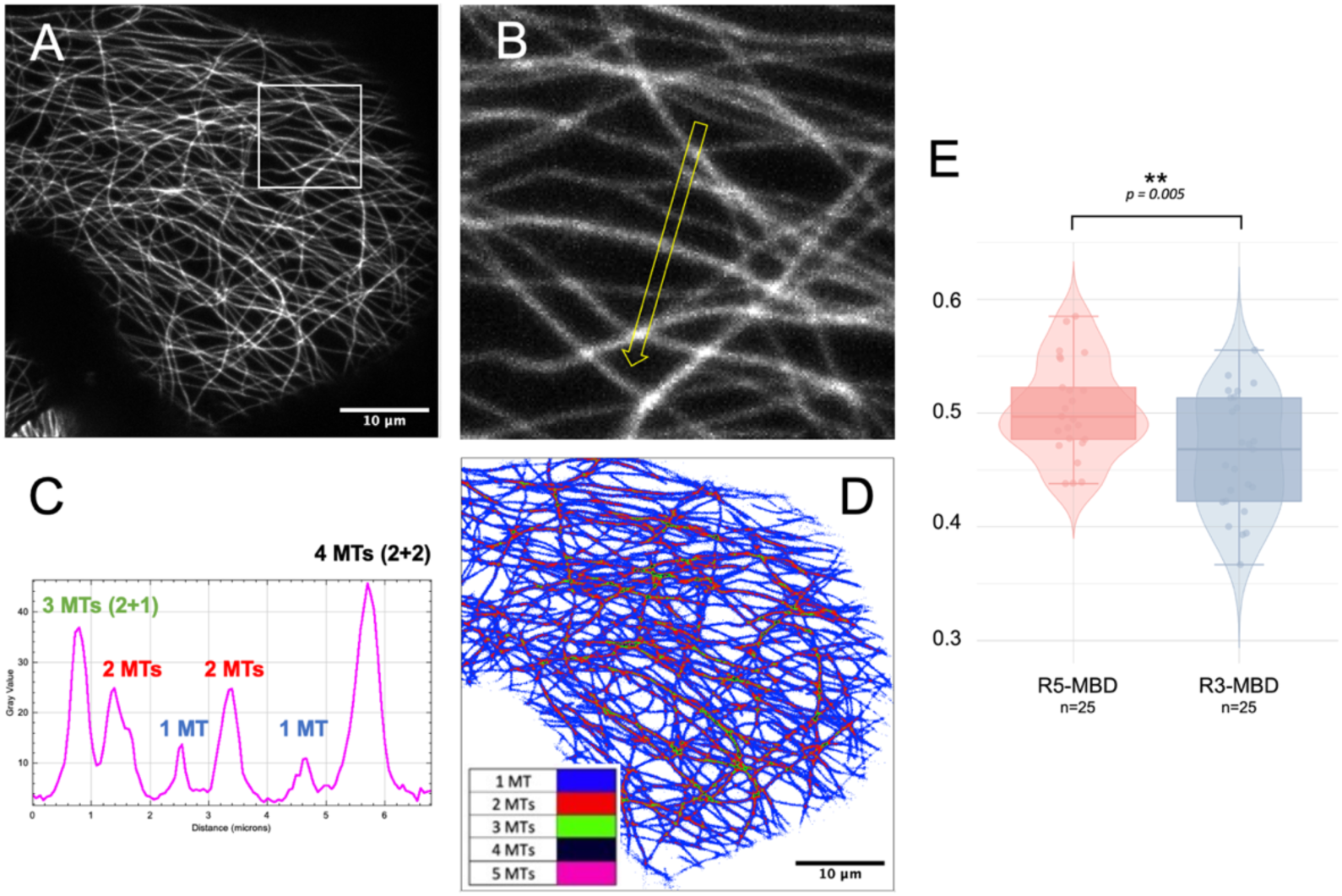
Bundling analysis of the 3R-MBD marker. **(A)** An *Arabidopsis* cotyledon cell expressing an mCitrine-3R-MBD fusion under the control of the UB10 promoter (maximum projection of a 2.5 µm-deep 3D stack). **(B)** Closeup (boxed in **(A)**) and **(C)** profile plot of the line shown in **(B)**. This shows that the MT marker signal, even in standard quality confocal images, is proportional to the number of underlying MTs, allowing quantification of MT bundles within the image. **(D)** The 8-bit image shown in **(A)** was divided by 14 and pseudo-colored. A gray value of 1 (blue) corresponds to single MTs, while gray values of 2 or 3 correspond to bundles of 2 or 3 microtubules respectively. Gray values of 0, 1, 2 or 3 in the divided image respectively correspond to gray level ranges of 0-6, 7-20, 21-34 and 35-48 in the original image. The addition of signal is easily seen at MT crossings, where two single MTs (gray value = 1, blue) crossings are colored in red (gray value = 2). **(E)** Box and violin plots showing the comparison of MT bundling rates between the original 5R construct and the new 3R one, upon expression in stably transformed *Arabidopsis* cotyledon cells.

From the stable *Arabidopsis* lines described in **Table 1**, we selected a 3R and a 5R line exhibiting comparable, medium expression levels of the MT marker, in order to compare their bundling activity. We then produced an image dataset of 25 epidermal cotyledon cells for each line. Using the imaging/analysis pipeline described above, we showed that MT bundling is significantly reduced with the 3R-MBD marker as compared to its 5R- counterpart (**Fig. 2E**). Although the difference is small, it is highly significant (t-test, p=0.005). Despite this improvement over the original 5R version, it remains that the 3R marker is also prone to causing developmental alterations in lines with the highest expression levels (data not shown). Therefore, standard precautions must apply for marker line selection, searching for an appropriate balance between brightness and absence of significant phenotype.

We finally defined a standard MT reporter as a PUB10::mCitrine::FS::3R-MBD::T_RbcsE9_ assembly (**Fig. 1B**). In our hands, this construct yields a bright and homogeneous MT signal and a tractable level of abnormal bundling or toxicity. In this construct, expression of the marker is driven by the *Arabidopsis* UB10 promoter [44–46], as MT markers driven by this promoter are considered far superior to 35S-driven ones, their expression being lower and more homogeneous in roots and aerial tissues [47]. The 3R-MBD was N-terminally fused to mCitrine through the short flexible linker (GlyGlyGlyGlySer), and flanked by an RbcsE9 terminator from *P. sativum* [48], a potent terminator that efficiently prevents transcriptional read-throughs and potential silencing issues.

### A three-color marker labeling the cell contours, nucleus and microtubules

We then incorporated the 3R-MBD microtubule marker into a triple marker assembly allowing to simultaneously label the cell contour, the nucleus and the microtubule cytoskeleton. Indeed, a cell contour marker often proves essential in many cell biology studies, especially for 3D approaches. Such a membrane or cell wall fluorescent marker allows segmentation of the cell volume, and quantitative assessment not only of several cell morphological parameters (shape and size), but also of the spatial distribution and organization of any internal component (vesicles, fibrils, etc.). A nuclear marker is also of high interest, as it enables visualization of the nucleus, its position and morphology, as well as the mitotic chromosomes and the metaphase plate. However, a main challenge here was to simultaneously optimize three independent markers for signal brightness, signal-to-noise ratio and spectral compatibility, while keeping cytotoxicity at a minimal level.

#### Assembly of a single construct for three fluorescent fusions

Among the wide variety of identified cell membrane markers, we set out to use the well-characterized RCI2a (At3g05880) that, in our hands, gives a reliable fluorescence signal without inducing cellular phenotypes when expressed from a moderate strength promoter such as the UB10 promoter. This 54-residue integral membrane protein was initially recovered from a general screen for markers of subcellular structures in *Arabidopsis* [49], and used as a membrane marker in several subsequent studies.

The nuclear marker was initially designed as a PCNA1-mTurquoise2 construct driven by its own promoter, allowing to monitor both nucleus morphology and cell-cycle progression [8]. However, in our hands, the fluorescence level and signal-to-noise ratio of the PCNA1- mTurquoise2 fusion did not prove sufficiently reliable for efficient segmentation of the nucleus, and we finally opted for a classical histone 2B marker. H2B.2 (*aka* HTB2, At5g22880) is a Group1 H2B protein peaking in S phase [50], that has been widely used as a nuclear marker since its original publication [51]. Here we chose to drive its expression by its endogenous promoter, in order to avoid the toxicity of non-assembled histones when expressed by non-native promoters (Frédéric Berger, personal communication). Despite H2B.2’s expression peak in S-phase, the H2B.2 promoter provides constant labeling of the nuclear DNA over the whole cell cycle in somatic tissues [50].

The final GoldenBraid assembly in a pDGB3-Omega destination vector (18 kb) contains all three markers (3R-MBD, RCI2A, and H2B.2) in fusion to mCitrine, tdTomato, and mTurquoise2 respectively, providing labelling of microtubules in yellow, membrane in red, and nucleus in cyan (**Sup Fig. 1**). A plant selection marker encoding resistance to Basta is also present in the T-DNA. Given the versatility of the GoldenBraid cloning system, colors can be easily swapped, or the selection marker changed if needed.

#### Transformation and selection of marker lines in *Arabidopsis* and *Capsella*

The pDGB3-MNM construct was used for *Agrobacterium*-mediated transformation of both *Arabidopsis* and *Capsella rubella*.

For *Arabidopsis*, about 100 primary transformants were recovered and assessed for fluorescent signal quality by confocal imaging. 41 lines displaying a consistent fluorescence signal for all three markers were selected for genetic analyses, among which 9 segregating for a single insertion locus were further characterized. These remaining lines were carefully examined for any growth or development phenotypes, as well as for signal stability and absence of silencing across 3 generations. A final line was chosen on the basis of signal quality across all three channels and absence of detectable phenotype or silencing. This reference line was called MNM. Its genome was fully sequenced and compared to its progenitor Col0, revealing a single T-DNA insertion on chromosome 5 between positions 22991671 and 22991692. The insertion site lies in the vicinity of the promoter of the At5g56860 gene, harbors a 20 bp deletion and a 63 bp insertion of unknown origin (**Sup Fig. 2**).

For *Capsella*, about 40 primary transformants were recovered and assessed for fluorescent signal quality by confocal imaging. 15 lines displaying a fluorescence signal for all three markers were selected for genetic analyses, among which 5 segregating for a single insertion locus were further characterized. These remaining lines were carefully examined for any growth or development phenotypes, as well as for signal stability and absence of silencing across 3 generations.

#### Phenotypic analysis of the MNM line in *Arabidopsis*

We set out to compare the developmental phenotype of the MNM line with its wild-type progenitor Col0. At the macroscopic level, root architecture of the homozygote MNM was undistinguishable from the wild-type (**Fig. 3A**). Root growth was then assessed *in vitro* on vertical agar plates. Growth between day 2 and day 7 after germination did not differ significantly between the two genotypes (**Fig. 3B**). At the subcellular level, we used calcofluor staining and confocal imaging to reveal cellular organization and track any defect in cell division plane orientation and cell morphology. Visual inspection and counting of cell files did not reveal any defects in the MNM line (**Sup Fig. 3**).

To characterize rosette growth and development, we compared wild-type Col-0 plants with homozygous MNM plants, and heterozygous F1 plants obtained by backcrossing a homozygous MNM line to Col-0. Plants were grown in the greenhouse and imaged from day 11 to day 25 after sowing (**Fig. 3C**). Analysis of the projected area of the rosette from D11 to D25 (**Fig. 3D**) showed that the heterozygous MNM line did not differ from the wild-type Col0 under these conditions. In contrast, the homozygous MNM line displayed a slight reduction in growth, especially at the D20-D22 stage, when the relative growth rate is maximal in the wild-type.

Globally, our phenotyping results show that the MNM line has very little root phenotype, if any. As for the aerial parts, we observed a slight growth reduction at late stages of rosette development in the homozygous MNM, but not in the heterozygote. As noted before, all cellular fluorescent markers display at least some level of toxicity, and as expected the combination of three co-expressed markers in the MNM line is no exception. However, the level of toxicity is minimal, and no phenotype is detectable at the heterozygous state in our conditions.

#### Validation of the 3-color marker in *Arabidopsis* and *Capsella*

The 3-color lines were subjected to confocal imaging on various confocal setups, including the Zeiss Spinning Disk, LSM710 and LSM980 Airyscan, and Leica SP5, SP8 and Stellaris systems. Examples of images obtained for various tissues of *Arabidopsis* are shown in **Fig. 4 and 5**. All images are pseudo-colored in the CMY format, where the microtubule/mCitrine channel is in yellow, the membrane/tdTomato in magenta, and the nucleus/mTurquoise in cyan.

**Figure 3.**
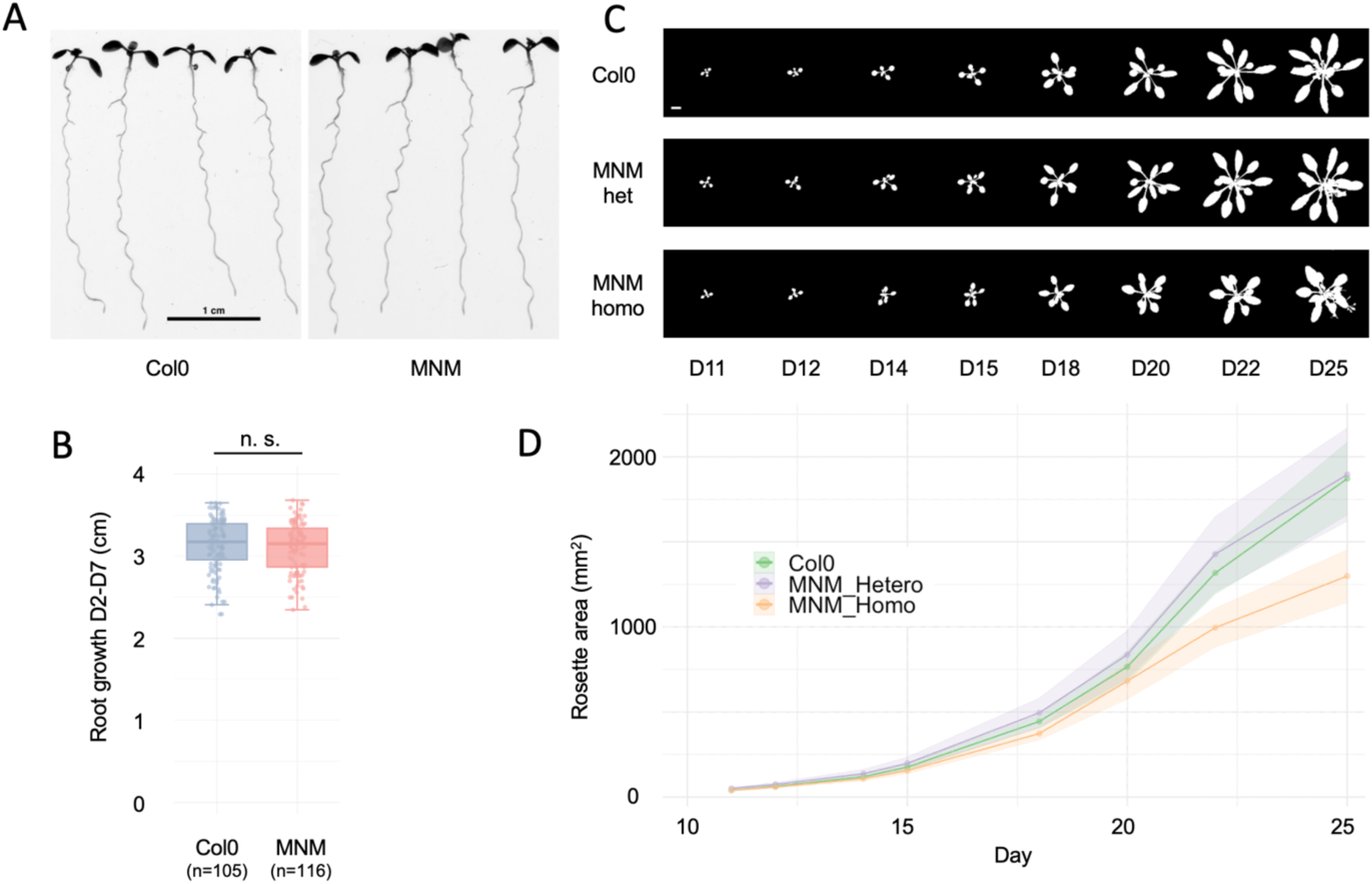
Phenotypic analysis of the *Arabidopsis* MNM line. **(A)** Root growth. Col0 and homozygote MNM plantlets were grown *in vitro* on vertical agar plates for each genotype. Root length was scored at days D2 and D7. (B) Root growth between D2 and D7 is not significantly different between the wild-type Col0 and the MNM homozygous line (Mann-Whitney, p = 0.35). **(C)** Rosette growth. Col0, homozygote and heterozygote MNM plants (14 of each) were grown in the greenhouse in standard conditions. Projected rosette surface was scored from day D11 to day D25 after sowing. **(D)** Growth curve from D11 to D25. The heterozygous MNM line (purple) is not different from the wild-type (green), but the homozygote MNM line (orange) displays a clear growth retardation, especially at the D20-D22 stage, where the daily relative growth rate (RGR) drops to 23%, compared to 36% in the wild-type background (Mann-Whitney, p = 0.007) and 35% in the heterozygous MNM background. Between D22 and D25, the daily RGRs amount to 14 % for the wild-type and 10 % for the homozygote (p = 0.003), whereas the heterozygote is at 11 % (p = 0.05032).

**Figure 4.**
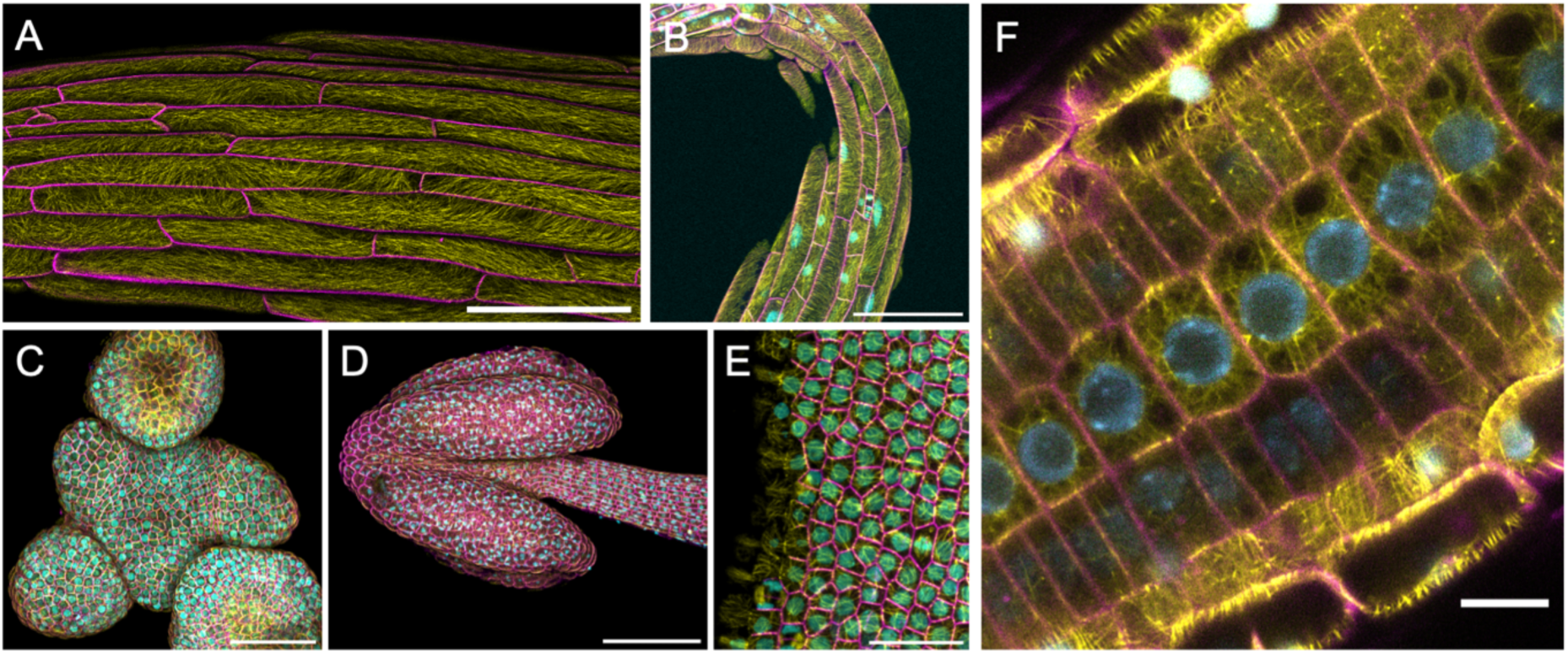
Various organs imaged from the *Arabidopsis* MNM line. **(A)** Two-channel (mCitrine + tdTomato) image of hypocotyl cells. Scale bar is 100 µm. **(B)** 3- color image of an apical hypocotyl cross from an etiolated plantlet. Scale bar is 100 µm. **(C)** 3- color image of a wild-type shoot apical meristem. Scale bar is 50 µm. **(D)** 3-color image of a wild type anther. Scale bar is 100 µm. **(E)** 3-color image of petal epidermal cells. Scale bar is 20 µm. **(F)** Root image obtained on a Zeiss LSM980 AiryScan microscope, revealing endoplasmic microtubules in epidermal cells. Scale bar is 10 µm.

**Figure 5.**
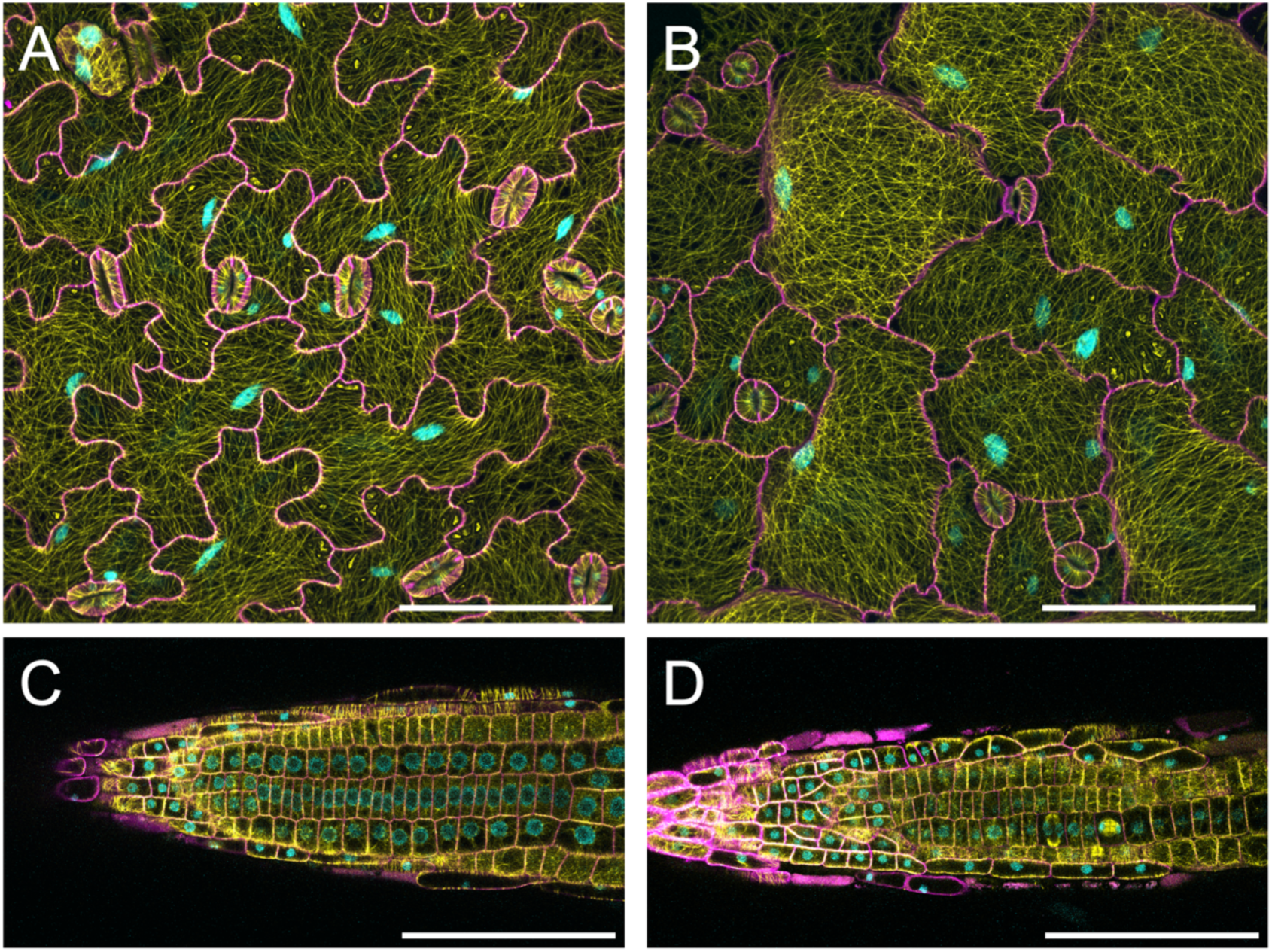
Mutant analysis with the MNM marker. Comparison of the wild-type *Arabidopsis* Col0 **(A, C)** with the *botero* mutant **(B, D)** using the MNM triple marker. **(A)** Wild-type *Arabidopsis* cotyledon epidermis. Scale bar is 100 µm. **(B)** *botero Arabidopsis* cotyledon epidermis. Scale bar is 100 µm. **(C)** Wild-type *Arabidopsis* root. Scale bar is 100 µm. **(D)** *botero Arabidopsis* root. Scale bar is 100 µm.

The *Arabidopsis* MNM line was tested by imaging a variety of organs and developmental stages. Expression of all three markers was detected in all tested organs including vegetative (root, hypocotyl, cotyledons, shoot apical meristem) and reproductive tissues (anther, fruit) (**Fig. 4**). It is worth noting, however, that as the H2B.2 promoter is not active in female gametophytes [50], the histone marker is most probably not expressed in these cells. Altogether, the three markers are expressed throughout the plant and are amenable to high-resolution confocal imaging from the subcellular to the whole organ level, showcasing the versatility and reliability of the triple marker. Although expectedly less bright than its homozygote counterpart, the heterozygous MNM line still provided images with a high signal-to-noise ratio.

In the root, the MNM line was imaged on a high-resolution AiryScan Zeiss confocal setup (**Fig. 4F**), revealing subtle details including well resolved endoplasmic microtubules. Furthermore, the MNM line was used for long-term live imaging of the growing root tip according to [52]. Images were acquired every 5 minutes for up to 12 hours (**Supplemental Movie**). All major microtubule arrays associated with the cell cycle were clearly detected, including the preprophase band (PPB), mitotic spindle, phragmoplast, and interphase cortical microtubule arrays, as well as the centromere within the nucleus. Notably, the developing cell plate was initially not labeled and became visible only after fusion with the plasma membrane of the mother cell. Photobleaching of the signal was present in all channels, but at a low level compatible with long-term imaging. This confirmed the potential of the MNM line for live imaging studies.

We then used the MNM marker to analyze phenotypic differences between wild-type *Arabidopsis* and a classical cytoskeleton/developmental mutant of *Arabidopsis*, *botero1-6* [28,53–55]. *Botero1-6* is a katanin mutant affecting microtubule severing and dynamics, with strong cellular and developmental defects from the cellular to the whole plant level (**Fig. 5**). As expected, comparison of the two genotypes using the MNM markers revealed strong differences in terms of cellular morphology and microtubule array patterns, in the cotyledon epidermis or in the root tip (**Fig. 5**).

Finally, successful introduction of the MNM marker in *Capsella rubella* (**Fig. 6**) showed that the MNM marker, originally optimized in *Arabidopsis*, can also be used in other plant species.

**Figure 6.**
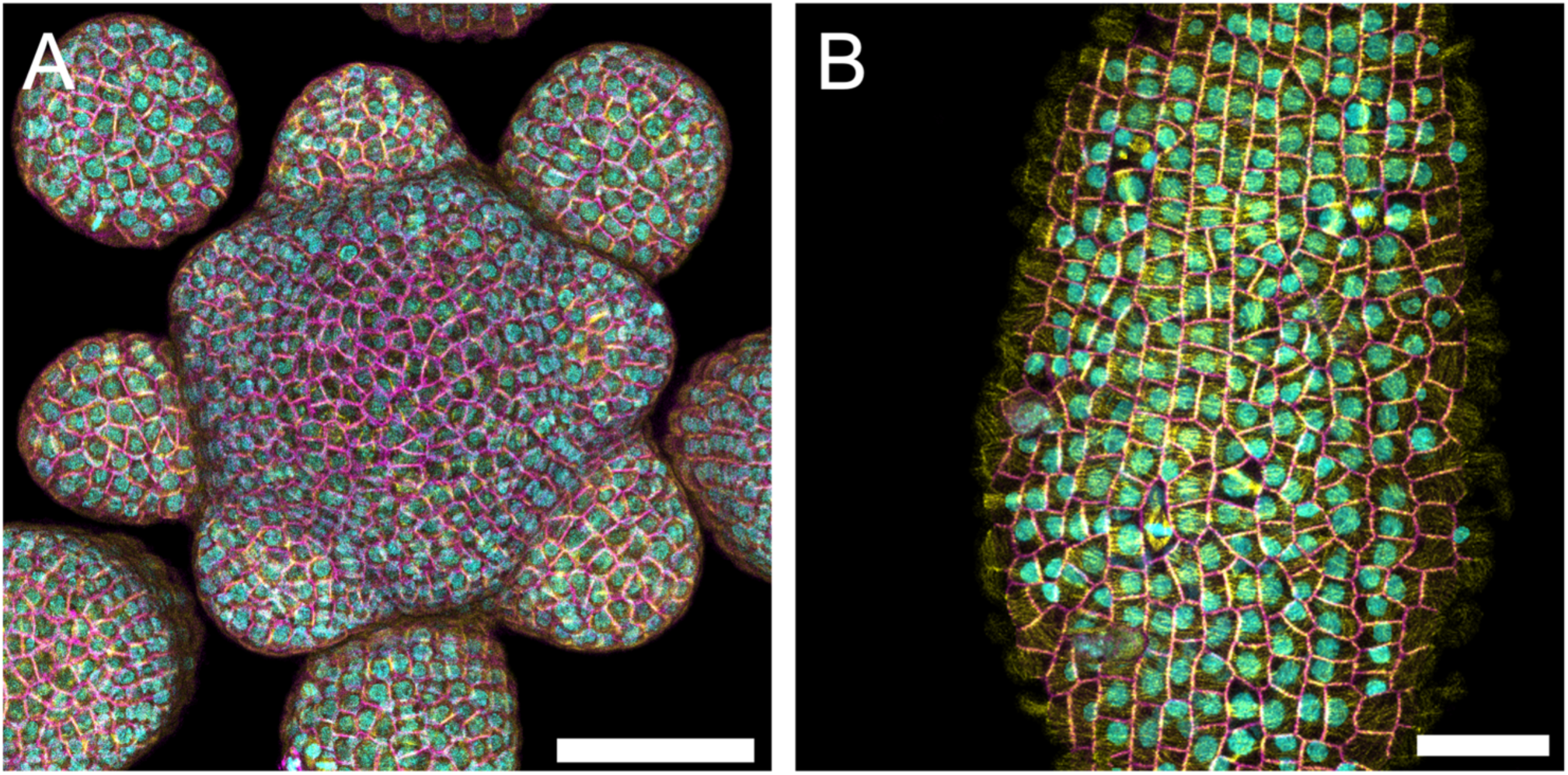
A *Capsella rubella* MNM line. **(A)** Shoot apical meristem of *Capsella*. Scale bar is 50 µm. **(B)** Early gynoecium at the 8 DAI (days after initiation) stage. Scale bar is 25 µm.

## Conclusion

Altogether, our results show that the single-locus triple MNM marker provides crisp confocal images of the cell contour, nucleus and microtubule cytoskeleton in various organs and cell types, in various imaging setups and in two plant species. The use of medium-strength promoters to drive the expression of subcellular markers facilitates the isolation of lines mostly devoid of significant developmental phenotypes, while maintaining a high level of fluorescence for all three markers even in the heterozygous state. Although the precise level of bundling and toxicity induced by the new 3R-MBD marker remains to be quantified with more precision in plant cells, they seem lower than that of the widely used 5R version, without compromising the quality of MT cytoskeleton labeling. The MNM marker holds great potential for 3-color imaging of plant development and morphogenesis at various scales, from the subcellular to the tissular levels.

## Supplementary data

**Supplemental Figure 1.** Graphical map of the pDGB3-MNM vector

**Supplemental Figure 2**. T-DNA insertion in the MNM line

**Supplemental Figure 3.** Absence of significant root phenotype in the MNM line

**Supplemental File 1:** Oligonucleotides and templates used to clone all GoldenBraid parts

**Supplemental File 2:** Transcription units used in this paper

**Supplemental File 3:** Full DNA sequence of the MNM vector (SnapGene format)

**Supplemental File 4:** ImageJ macro for bundle analysis

**Supplemental File 5:** User guide for ImageJ macro

**Supplemental Movie 1:** Time-lapse imaging of the root apex of the *Arabidopsis* MNM line. The same root was acquired every 5 minutes for 12 hours as a 3D confocal stack (16 slices). Here a single plane (slice 5) is shown.The scale bar is 10 µm. The time is indicated as hours:minutes. The microtubule/mCitrine channel is pseudo-colored in yellow, the membrane/tdTomato in magenta, and the nucleus/mTurquoise in cyan.

## Acknowledgments

The authors are grateful to Frédéric Berger for helpful discussions on the choice of the histone marker, to Elliot Meyerowitz for the gift of a RCI2A clone, and to Delphine Charif for performing the alignment of sequencing reads against the reference genome. This work has benefited from the support of IJPB’s Plant Observatory platforms, PO-Plants and PO-Cyto, and from Saclay Plant Sciences support in the frame of "Educational Projects" and "Shared Tools" calls.

## Data availability

The data that support the findings of this study are available from the corresponding author upon request.

---

**Supplemental Figure 1.**
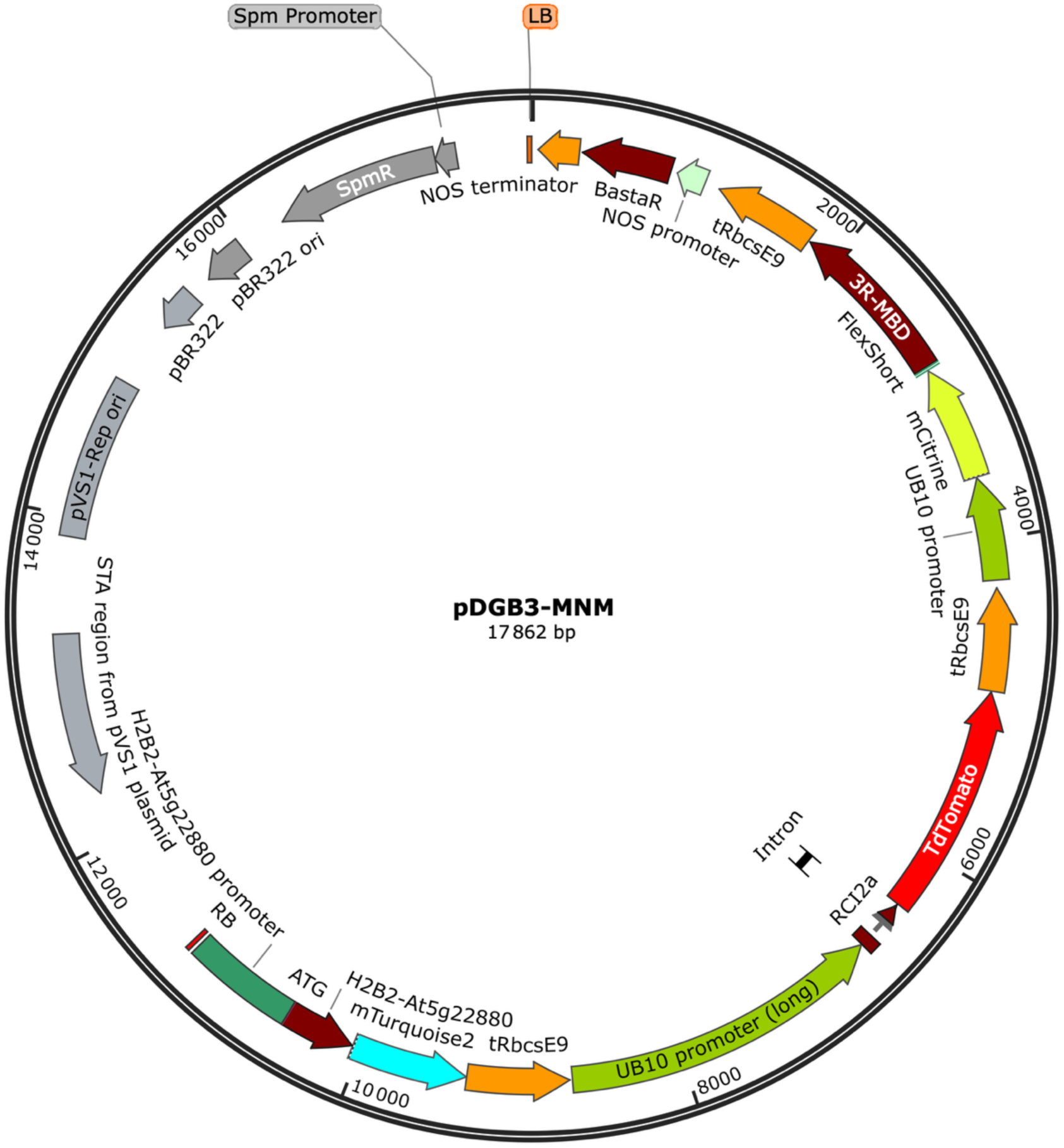
Graphical map of the pDGB3-MNM vector. The vector is in a pDGB3-Omega1 backbone. The 3R-MBD and RCI2A markers are driven by UB10 promoters, the H2B.2 is driven by its own promoter, and all three markers are flanked in 3’ by the RbcsE9 terminator. The Basta selection marker is flanked by the NOS promoter and terminator. The three fluorescent fusions and the Basta selection marker are all in the same orientation on the T-DNA, in order to avoid the production of antisense RNAs that may induce silencing.

**Supplemental Figure 2.**
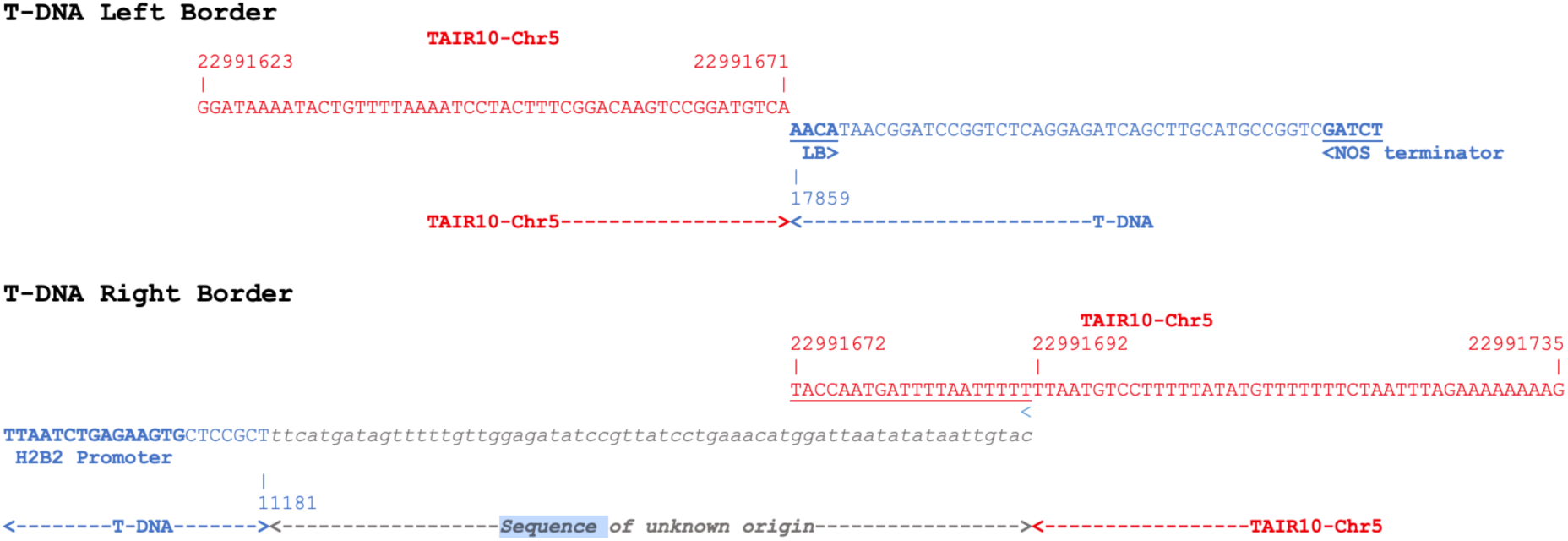
T-DNA insertion in the MNM line. A single T-DNA from the pDGB3-MNM vector is present in the MNM line, inserted on chromosome 5 between positions 22991671 and 22991692 (TAIR10 coordinates), thus corresponding to a 20 bp deletion (underlined), together with a 63 bp insertion of unknown origin. Sequence coordinates of the T-DNA refer to the pDGB3-MNM sequence (Supplemental File 3). The *Arabidopsis* chromosome 5 is in red, the T-DNA in blue and the insertion sequence in gray.

**Supplemental Figure 3.**
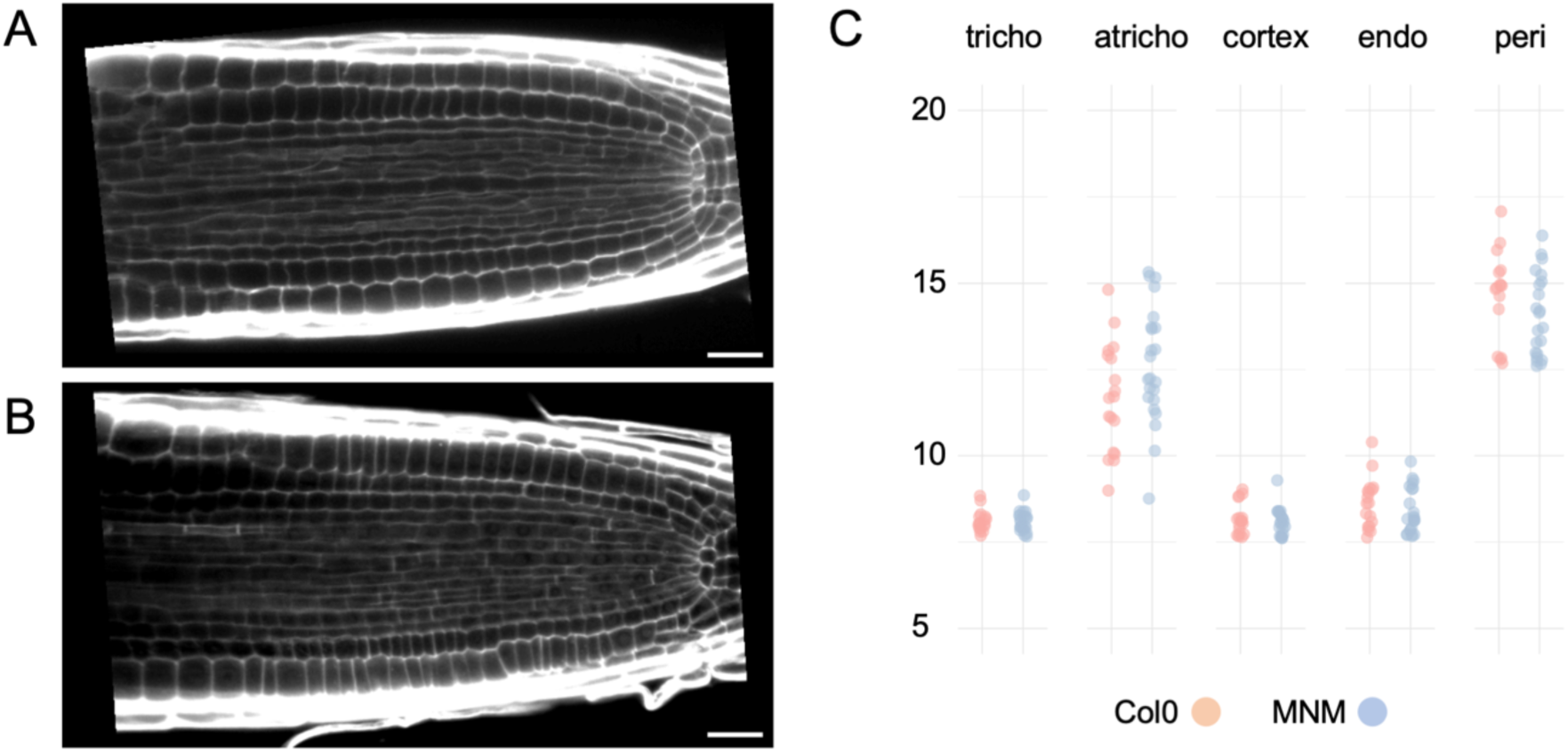
Absence of significant root phenotype in the MNM line. Representative examples of a median longitudinal section from a Col0 **(A)** and MNM root **(B)** stained with Calcofluor and acquired as a 3D confocal stack. Scale bars are 20 µm. A total of 6 roots were acquired for each genotype. Visual inspection of the stacks did not reveal any defects in MNM roots as compared to the wild-type, in terms of cellular organization, cell shape or size. Cell files **(C)** were scored at various distances from the Quiescent Center (50, 100, 150 and 200 µm). For all tissues, the number of cell files did not differ significantly between genotypes (Mann and Whitney, α = 0.05).

