## Supplemental File 4 for "A triple fluorescent marker for live imaging of plant cell morphogenesis": Supplemental File 4.ijm

```
// ImageJ macro for bundle analysis 
// The input directory must contain a sub-directory named "INPUT" containing the original stacks
// An output directory named "OUTPUT" will be created by the macro


#@ File (label = "Input directory", style = "directory") inputdir
#@ String (label = "File suffix", value = ".tif") suffix

input = inputdir + File.separator + "INPUT";
output = inputdir + File.separator + "OUTPUT";
File.makeDirectory(inputdir + File.separator + "OUTPUT");

setBackgroundColor(0, 0, 0);
setForegroundColor(255, 255, 255);

processFolder(input);


selectWindow("Log");
	saveAs("text", output + File.separator + "Log.xt");
	run("Close");


// function to scan folders/subfolders/files to find files

function processFolder(input) {
	list = getFileList(input);
	list = Array.sort(list);
	for (i = 0; i < list.length; i++) {
		if(File.isDirectory(input + File.separator + list[i])) {
			processFolder(input + File.separator + list[i]);
		}
	else {
		if(endsWith(list[i], suffix))
		processFile(input, list[i]);
	}
	}
}

function processFile(input, file) {

open(input + File.separator + file);
Name = File.nameWithoutExtension;


// Make a MAX proj of the original Stack and save it in the output folder
selectImage(file);
rename("Stack");
setSlice(8);
run("Enhance Contrast", "saturated=0.35");

run("Z Project...", "projection=[Max Intensity]");
run("Enhance Contrast", "saturated=0.35");
saveAs("Tiff", output + File.separator + Name + "_Max.tif");
rename("Max-proj");


// Make a mask to remove background between MTs:
// Apply a median filter (1 pixel), erode, dilate, binarize and divide by 255

// Start by running a 3D median filter on the original Stack
selectImage("Stack");
run("Mean 3D...", "x=1 y=1 z=1");

// Then make a Z-max-proj of the resulting stack and make a mask
run("Z Project...", "projection=[Max Intensity]");
setOption("BlackBackground", true);
run("Convert to Mask");
run("Erode");
run("Dilate");
rename("Mask1");
run("Duplicate...", "title=Mask");
selectImage("Mask1");
run("Divide...", "value=255");
saveAs("Tiff", output + File.separator + Name + "_Mask.tif");
rename("Mask1");
imageCalculator("Multiply", "Mask1","Max-proj");
selectImage("Mask1");
// this is now the result of the division
run("Grays");
run("Enhance Contrast", "saturated=0.35");
saveAs("Tiff", output + File.separator + Name + "_Max-Masked.tif");
rename("Max-Masked");


run("Tile");

// Define region of interest
selectImage("Mask");
setTool("freehand");
waitForUser("Select the region of interest in the Mask, then click OK to continue");

// Transfer selection to the Max-Masked image and crop
selectImage("Max-Masked");
run("Restore Selection");
run("Clear Outside");
saveAs("Tiff", output + File.separator + Name + "_Max-Masked-Cropped.tif");
rename("Max-Masked-Cropped");


close("Mask");
close("Stack");

// Tile the "Max-proj" and the "Max-Masked-Cropped" images
run("Tile");

// Analyze MT grey values
selectImage("Max-Masked-Cropped");
getRawStatistics(nPixels, mean, min, max, std);
run("Histogram");
rename("Histogram");
Table.showHistogramTable;

// Make line profiles to help estimate single MT grey value

selectImage("Max-Masked-Cropped");
run("Select None");
setTool("line");
run("Line Width...", "line=3");
xm = getWidth()/2;
ym = getHeight()/2;
makeLine(xm-50,ym,xm+50,ym);
run("Plot Profile");
Plot.enableLive(true);
Plot.setStyle(0, "blue,none,3.0,Line");

waitForUser("Draw a line across several single MTs\nin order to estimate the average grey value\nInspect histogram table to define the grey of a single MT\nFrom these choose a division value");

// Clean desktop
close("Plot of Max-Masked-Cropped");
close("Histogram");
close("Stack");

selectImage("Max-Masked-Cropped");
run("Select None");
run("Duplicate...", "title=REF");
run("Out [-]");
run ("Tile");
selectImage("Max-Masked-Cropped");
run("glasbey");
run("In [+]");
run("In [+]");

waitForUser("On the next window, activate Preview, and test different division values.\nClick OK when ready");

run("Divide...");

close("Max-Masked-Cropped");


// Choose parameters
Dialog.create("Division option");
Dialog.addMessage("Choose division factor");
Dialog.addNumber("Choose division factor", 10);
Dialog.show();
div = Dialog.getNumber();

// do stuff

selectImage("REF");
run("Divide...", "value=&div");
rename("Glasbey");


// Saving glasbey image
selectImage("Glasbey");
getHistogram(hval, hcounts, 256);
resetMinAndMax;
saveAs("Tiff", output + File.separator + Name + "_Max_Masked_Cropped_Div.tif");

// Now for the calculations

MT01 = hcounts[1];
MT02 = hcounts[2];
MT03 = hcounts[3];
MT04 = hcounts[4];
MT05 = hcounts[5];


MTtot = MT01 + (2*MT02) + (3*MT03) + (4*MT04) + (5*MT05);
bundling = 1 - (MT01/MTtot);

print("Image:\t" + file);
print("MAX-Proj Selection Area (pixels):\t" + nPixels);
print("MAX-Proj Selection mean min max std:\t" + mean,min,max,std);
print("Histogram of divided image:");
Array.print(hcounts);
print("Division value:\t" + div);
print("Bundling rate:\t" + bundling);
print("------------------------------");
print("");

close("*");

}
```
