## Supplemental File 5 for "A triple fluorescent marker for live imaging of plant cell morphogenesis"

### Supplemental File 5. User guide for the ImageJ macro for bundle analysis.

#### Image acquisition recommendations

All 3D stacks must be acquired ensuring that no pixel was saturated. All 3D stacks were acquired with a voxel size of 60 µm x 60 µm x 170 µm.

#### Installing Fiji

Here is the link to download the Fiji software (Schindelin *et al.*, 2012):
https://imagej.net/software/fiji/downloads

#### Available images

Raw confocal images used for implementing and testing the macro are available upon request.

#### Starting files

The input directory must contain a sub-directory named "INPUT" containing the original Z-stacks. An output directory named "OUTPUT" will be created by the macro.
The Macro opens up four images:

The original Z-stack,

The Maximum Projection of the Z-stack,

The Mask of the microtubules (MT) signal,

The Mask applied to the Maximum Projection. Applying a mask to the maximum projection image allows measurement of signal values corresponding to MTs, while eliminating background noise from the analysis.

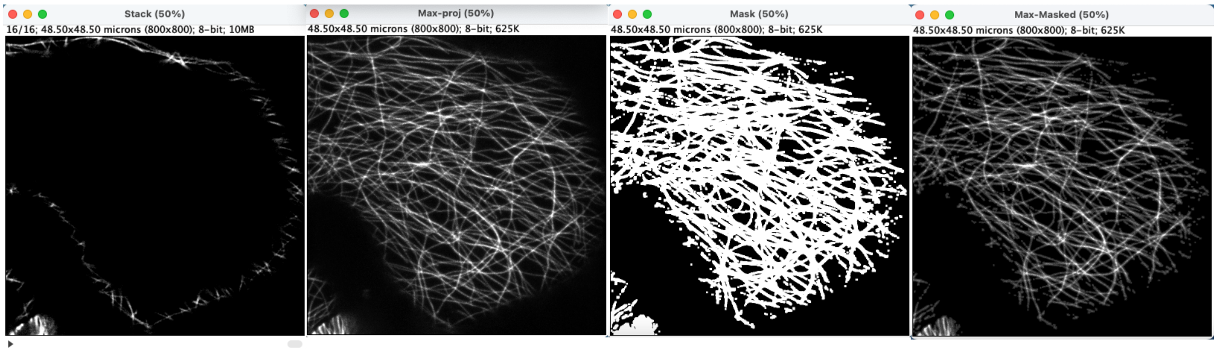


#### Drawing the ROI

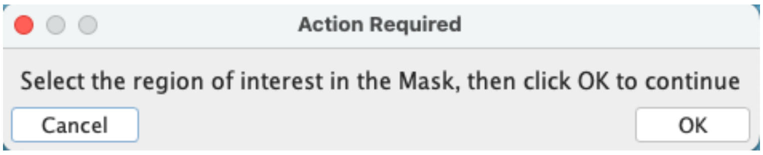


Browsing through the slices of the original Z-stack allows identification of an appropriate ROI for the next step. The selected ROI should correspond to a region where the signal is consistently present, ideally within the flattest area of the projection, while avoiding regions where MTs are only partially captured (i.e., not imaged across their full thickness).
Draw the ROI using the "Freehand selections" tool (here in yellow).

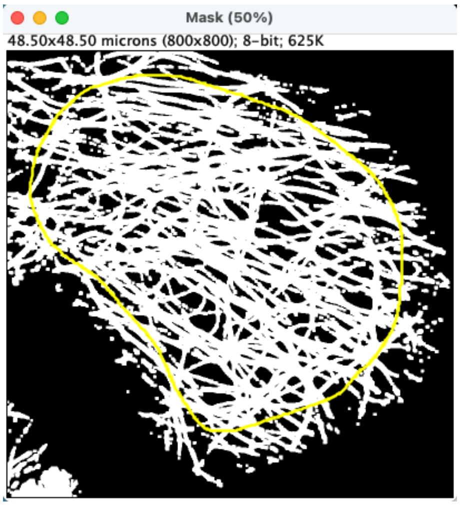


#### Selection of the mean value of the signal of single MTs

The macro then opens several windows to assist in selecting the mean signal value of a single MT. It allows the user to draw a line (shown in yellow below) across MTs in the maximum projection after application of the binary mask, and generates a live intensity plot profile to evaluate the signal of individual MTs.
It also opens a histogram along with its corresponding table, listing the counts for all gray values in the image. The first peak corresponds to the mean signal of single MTs (here 10).

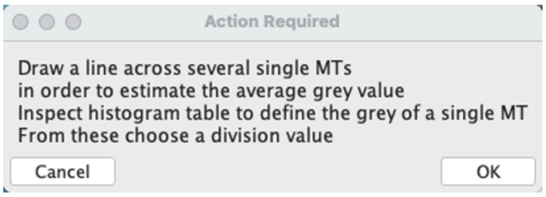


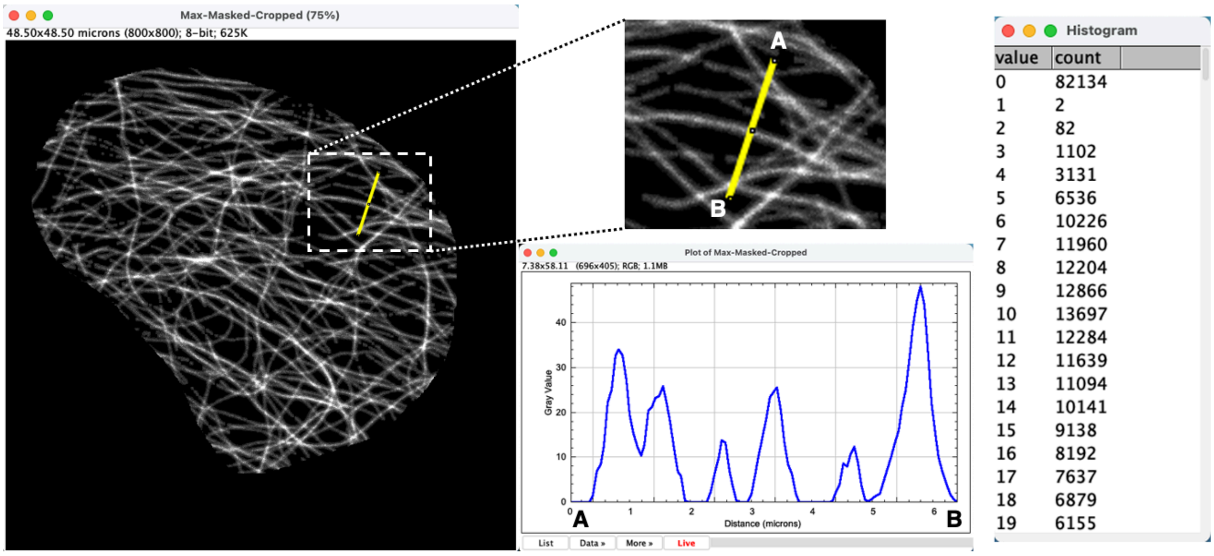


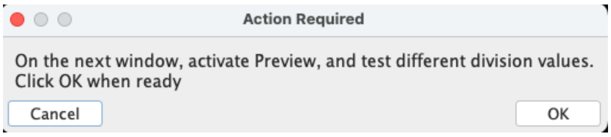

The macro opens then the mask-masked and cropped maximum projection with a "Glasbey" on dark LUT.

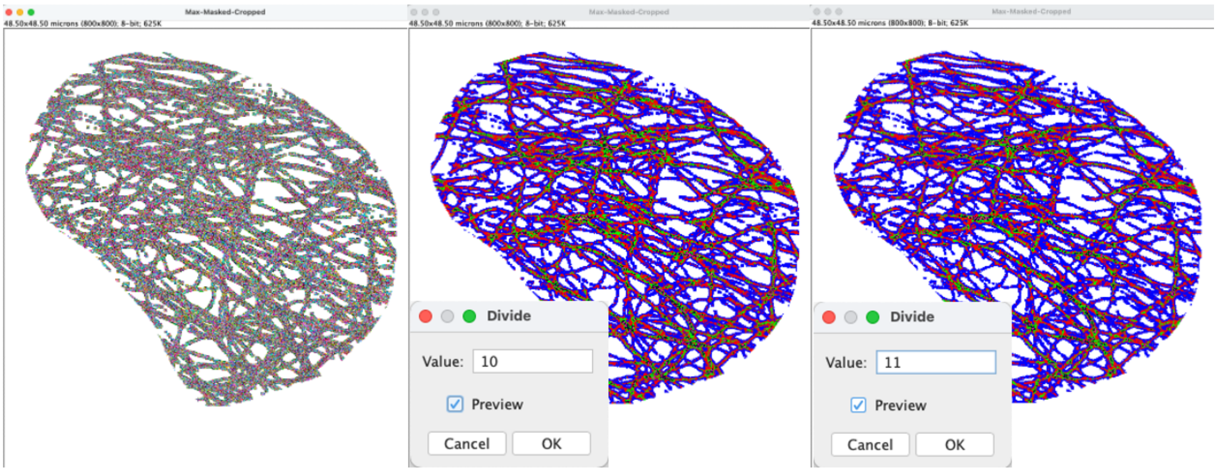


The Macro allows testing of different division values. Dividing the image by the value corresponding to single MTs enables the identification of individual MTs (pixels of value 1, here in blue), bundles of two MTs (pixels of value 2, pink), three (pixels of value 3, green), four (pixels of value 4, black), and so on.

Here we systematically used the value of the first peak, corresponding to the mean intensity of individual MTs, plus one (which equals eleven in the example shown). This ensures that most single MTs have a pixel value of 1 (blue).
Relying on histogram-derived values to analyze all images, rather than visual inspection of Glasbey LUT–segmented images, ensures an unbiased determination of the division factor across the entire image set.

When "OK" is activated in the "Divide" window, another window opens to choose the division factor (see below).

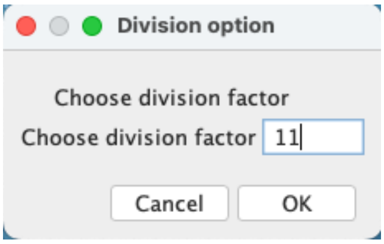


Simple pixel counting then allows to approximate the MT bundling rate (BR) by the formula:

MTtot = MT1 + (2*MT2) + (3*MT3) + (4*MT4) + (5*MT5)

BR = 1 - (MT1/MTtot)

The Macro then saves the log containing all the measured values and the calculated MT bundling rate.
